# Antigen presentation by clonally diverse CXCR5+ B cells to CD4 and CD8 T cells is associated with durable response to immune checkpoint inhibitors

**DOI:** 10.1101/2022.12.08.519660

**Authors:** Lizhong Ding, Lu Sun, Melissa T. Bu, Yanjun Zhang, Lauren N. Scott, Robert M. Prins, Maureen A. Su, Melissa G. Lechner, Willy Hugo

## Abstract

The mechanism by which ICI (immune checkpoint inhibitor) induce durable antitumor T cell activity remains inadequately defined. Tumors from melanoma patients who responded to ICI or MAPK pathway inhibitors (MAPKi) therapy generally displayed increased T cell infiltration and interferon gamma (IFNγ) pathway activation. Yet, the rate of durable tumor control after ICI is almost twice that of MAPKi. Comparing the transcriptome of cohorts of melanoma patients treated with ICI or MAPKi therapy, we discovered that response to ICI is associated with CXCL13-driven recruitment of CXCR5+ B cells with higher clonal diversity than MAPKi. Higher B cell receptor (BCR) diversity allows presentation of diverse tumor antigens by B cells, resulting in robust increases of IFNγ pathway activity and CXCL13 expression in tumor reactive CD8 T cells after ICI therapy. Accordingly, ICI-treated melanoma patients, but not MAPKi, whose tumors displayed higher BCR diversity and IFNγ pathway score, survived significantly longer than those with either one or none. Thus, response to ICI, but not to MAPKi therapy, induces the recruitment of clonally diverse antigen presenting B cells that activate tumor specific, cytotoxic CD8 T cells to effect a durable antitumor immune response. Our result suggests that enhancing B cells’ tumor antigen presentation to intratumoral CD8 T cells can increase the rate of long-term response to ICI therapy.

## Introduction

Metastatic melanoma used to have a dismal median overall survival of only nine months after diagnosis (*1*, *2*). However, the advent of immune and targeted therapies (*3*, *4*) has significantly improved the survival of patients with metastatic melanoma. The highly immunogenic nature of melanoma have made it the model cancer to study response to immune checkpoint inhibitors (ICI), such as the blocking antibodies against T cell inhibitory receptors (T cell “checkpoints”) such as cytotoxic T lymphocyte antigen (CTLA)-4 and programmed death protein (PD)-1 (*5*–*8*). Despite the remarkable success of ICIs in many patients with melanoma, their clinical response remains difficult to predict (*9*–*12*). Previous work highlighted the association of T cell infiltration and patient response to ICI; ICI-treated tumors displayed higher number of infiltrating T cells accompanied by expression of genes related to interferon pathway activities, demonstrating increased production of interferon gamma (IFNγ) of these T cells (*13*–*17*).

The discovery of constitutive activation of RAF/MEK/ERK signaling via the BRAF V600 mutation in nearly half of all cutaneous melanoma cases also revolutionized cancer therapy(*18*). MAPK pathway inhibitor (MAPKi) treatment significantly prolonged the survival of metastatic melanoma patients with BRAF V600 mutation (BRAF^V600^ mutant melanoma) (*19*–*21*). In addition to its direct tumor suppressive effect through MAPK pathway inhibition, MAPKi treatment also increases the infiltration of antitumor T cells into the tumor microenvironment (TME), suggesting immune modulatory effects (*22*, *23*).

Intriguingly, despite MAPKi therapy having a higher rate of initial response than ICI (MAPKi (dabrafenib+ trametinib): 67% (*19*), MAPKi (vemurafenib+cobimetinib): 68% (*20*) vs ICI (nivolumab and ipilimumab): 58% (*8*)), the 5-year survival rate of BRAF^V600^ mutant subset of melanoma patients treated with MAPKi is only around half that of ICI (34% using dabrafenib and trametinib (*24*), 31% using vemurafenib and cobimetinib (*25*) vs. 60% after nivolumab and ipilimumab (*26*)). Furthermore, a matching-adjusted study found that ICI treatment improved overall survival (OS) of BRAF-mutant melanoma patients when compared to MAPKi (*27*). While acknowledging some differences among these studies, this consistent and significant difference in durable survival between ICI and MAPKi treated melanoma patients suggests that there may be fundamental differences in the anti-tumor responses induced by these therapies.

To discover such differences, this study analyzes the changes in immune related gene expressions that are significantly associated with survival of melanoma patients after ICI or MAPKi treatment. Previous work compared on-treatment tumors (i.e., these tumor samples were biopsied after treatment) of patients responding to ICI (ICI OT-R) compared with those from patients not responding to ICI (ICI OT-NR) to nominate immune factors/pathways associated with response to ICI. However, this comparison is not necessarily informative since ICI OT-NR tumors generally have less immune infiltration compared to ICI OT-R hence most immune cell-related markers were upregulated in ICI OT-R group. Which of these are drivers of ICI’s durable response vs. insignificant bystanders is therefore unclear. Since MAPKi OT-R tumors also have more immune infiltration than MAPKi OT-NR tumors, yet MAPKi OT-R patients are less likely to achieve a durable response than ICI OT-R patients, we posit that immune genes/pathways that are upregulated/enriched in ICI OT-R, but not in MAPKi OT-R tumors, can help explain the higher rate of durable responses in ICI treated patients.

We discovered higher expression of genes related to B cells in the ICI OT-R tumor, which were accompanied by genes indicative of tertiary lymphoid structures (TLS) formation such as *CXCL13* and *CXCR5*. Single cell RNA-seq analysis of ICI-treated melanoma confirmed that response to ICI increased the number of germinal center-like B cells and its associated T follicular helper CD4 T cells, again, indicative of TLS formation as reported previously (*28*, *29*) (*30*, *31*). These cells were recruited into the TME by CXCL13-producing, cytotoxic CD8 T cell population that was specific to ICI. Importantly, BCR diversity, but not clonality, was significantly associated with extended overall survival after ICI but not MAPKi. The significant association between BCR diversity with survival after ICI suggests that ICI-induced B cells serve as antigen presenting cells, which will be able to present more diverse tumor antigens with more diversified BCR clones. Indeed, patients whose tumors display both higher BCR diversity and IFNγ signaling pathway score after ICI, which suggest successful antigen presentation by B cells to T cells, have significantly longer overall survival than those with either one or none. Our results suggest a combination of ICI with therapies that enhance antigen presentation by clonally diverse B cells can result in a more durable antitumor immune response.

## Results

### ICI and MAPKi treatment induce comparable levels of T cell infiltration

To perform a comparative analysis of transcriptomic response to ICI and MAPKi, we analyzed three separate immunotherapy datasets (*16*, *32*, *33*) and three targeted therapy datasets (*23*, *34*, *35*) (Supp. Fig 1A, Supp. Table 1A). All samples were classified into three groups: pre-treatment (PT), on-treatment responding (OT-R), and on-treatment non-responding (OT-NR). OT-R is defined by clinical benefit after therapy (CR, PR, or SD by RECIST criteria). OT-NR is defined by no clinical benefit (PD) (Supp. Table 1A). Gene expression in samples across the datasets were integrated and batch normalized (Supp. Table 1B, see Methods).

We asked whether the set of differentially expressed genes (DEG) between ICI and MAPKi could reveal any cell populations or biological processes that may explain ICI’s more durable antitumor response. To this end, we selected genes upregulated in the OT-R compared to the OT-NR samples for each therapy (Fig 1A, see Methods for details). DEG were grouped into either ICI-specific, MAPKi-specific, or ICI and MAPKi-shared (Fig 31B, Supp. Table 1C), and gene sets enriched by these DEGs were computed using Enrichr (*36*). We first noted gene sets related to T cells being enriched in the OT-R groups of both therapies (Fig 1C, Supp. Table 1D). The levels of general T cell marker (*CD3D*), cytotoxic CD8 T cells (*CD8B*) and T cell signature were significantly higher in the OT-R tumors of MAPKi or ICI compared to PT and OT-NR tumors (Fig 1D, Supp. Fig 1B, Supp. Table 1E).

**Fig 1:**
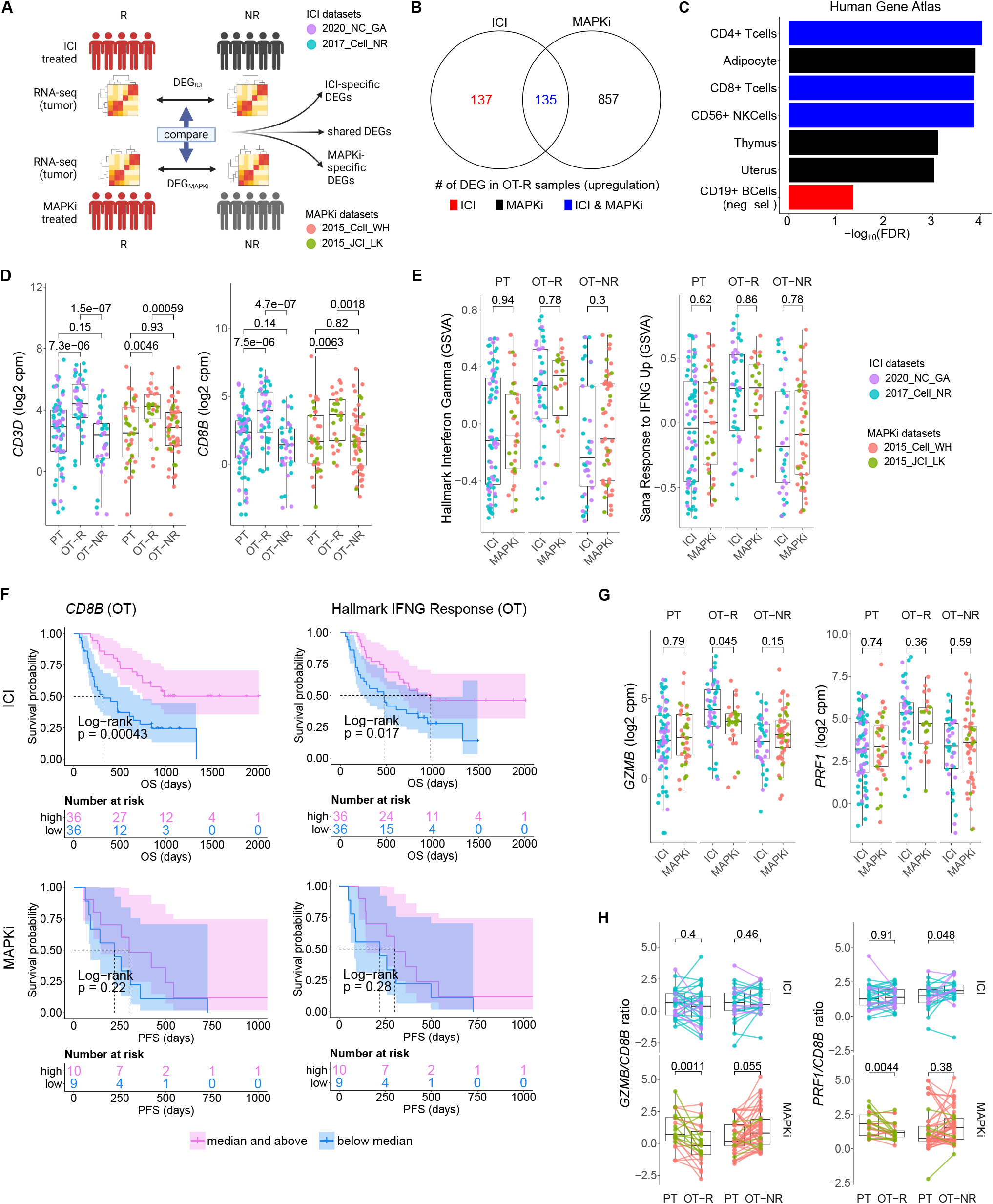
Response to ICI and MAPKi therapy are both associated with increased T cell infiltration and enhanced IFNγ pathway activity in the tumor. A) Schematic of comparative analysis between transcriptomic response to ICI and MAPKi therapy. B) The number of differentially upregulated genes in ICI OT-R (red), MAPKi OT-R (black) or both (blue). The differentially expressed genes (DEGs) are computed with respect to the OT-NR samples of each group. C) Enriched cell marker genes (based on Human Gene Atlas using Enrichr tool) in ICI-specific, MAPKi-specific, and ICI and MAPKi-shared DEGs in (B). D) Normalized expression of T cell marker genes *CD3D* and *CD8B* in the PT, OT-R and OT-NR samples of patients under ICI and MAPKi therapy. The dataset cohort of the datapoints is listed in the righmost legend. E) GSVA gene set enrichment scores of two interferon gamma downstream gene sets from the Molecular Signature database. Both ICI and MAPKi therapy induced a similar increase in the two pathways’ scores. F) Kaplan-Meier survival curves of ICI- or MAPKi-treated melanoma patients stratified by either *CD8B* expression (left) or hallmark interferon gamma response gene set scores (right) in their OT tumors. G, H) Normalized expression (G) and *CD8B*-normalized expression (H) of T cell cytotoxicity genes *GZMB* and *PRF1* in the PT, OT-R and OT-NR samples of patients under ICI and MAPKi therapy.

**Fig 2:**
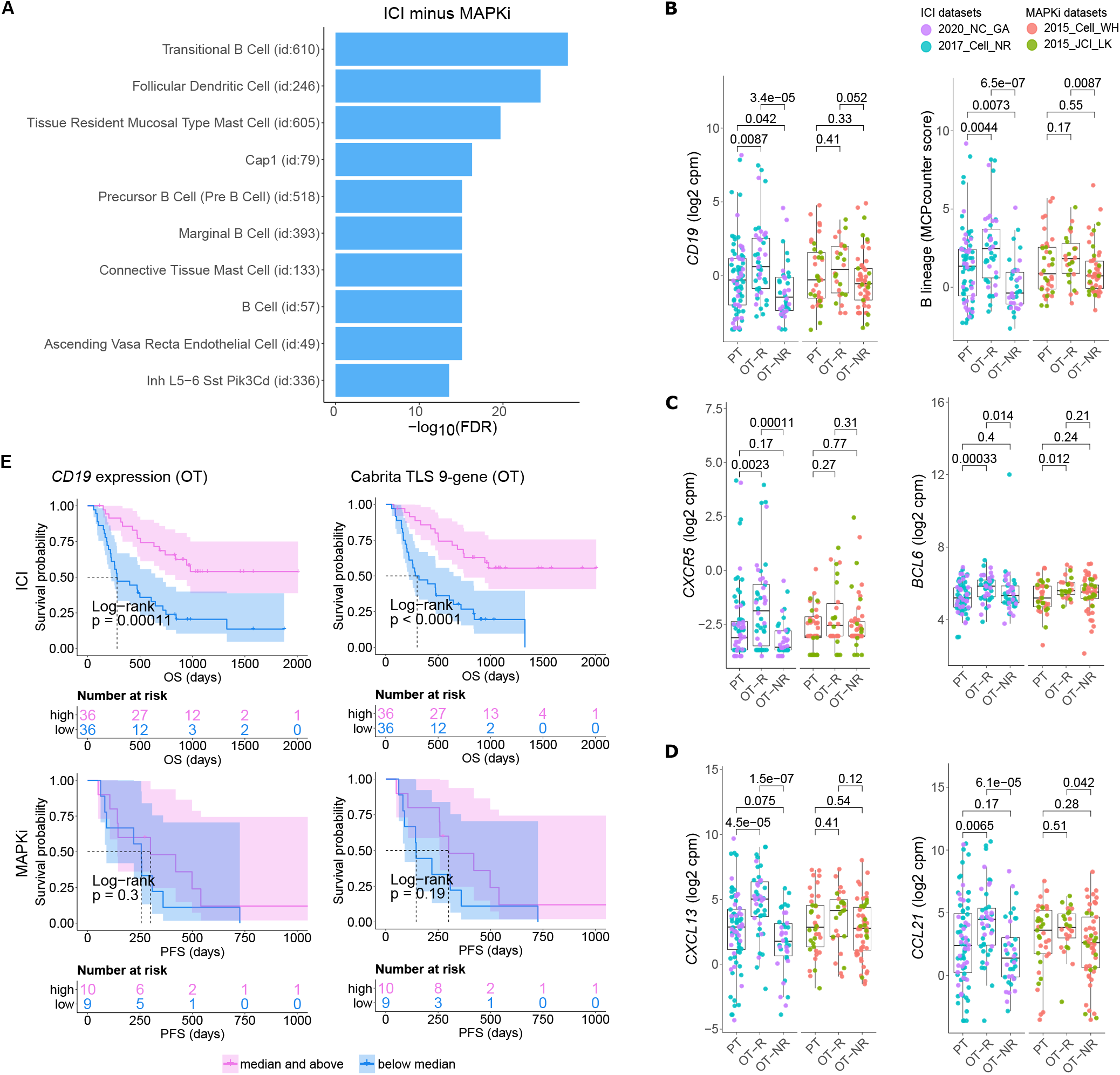
Relative increase of B cell and tertiary lymphoid structure (TLS) marker genes in response to ICI therapy compared to MAPKi. A) Enriched tissue specific gene sets in DEGs upregulated in ICI OT-R tumors with respect to MAPKi OT-R tumors (after adjustment by respective therapy group’s OT-NR tumors; see Methods). B) Normalized expression of B cell marker genes and enrichment of B cell lineage gene set among the PT, OT-R and OT-NR tumors in the ICI or MAPKi therapy group. C, D) Normalized expression of TLS-related genes, *CXCR5*, *BCL6*, *CXCL13* and *CCL21* among the PT, OT-R and OT-NR tumors in the ICI or MAPKi therapy group. E) Kaplan-Meier survival curves of ICI- or MAPKi-treated melanoma patients stratified by either *CD19* expression (left) or TLS gene set enrichment score (right) of their OT tumors.

**Fig 3:**
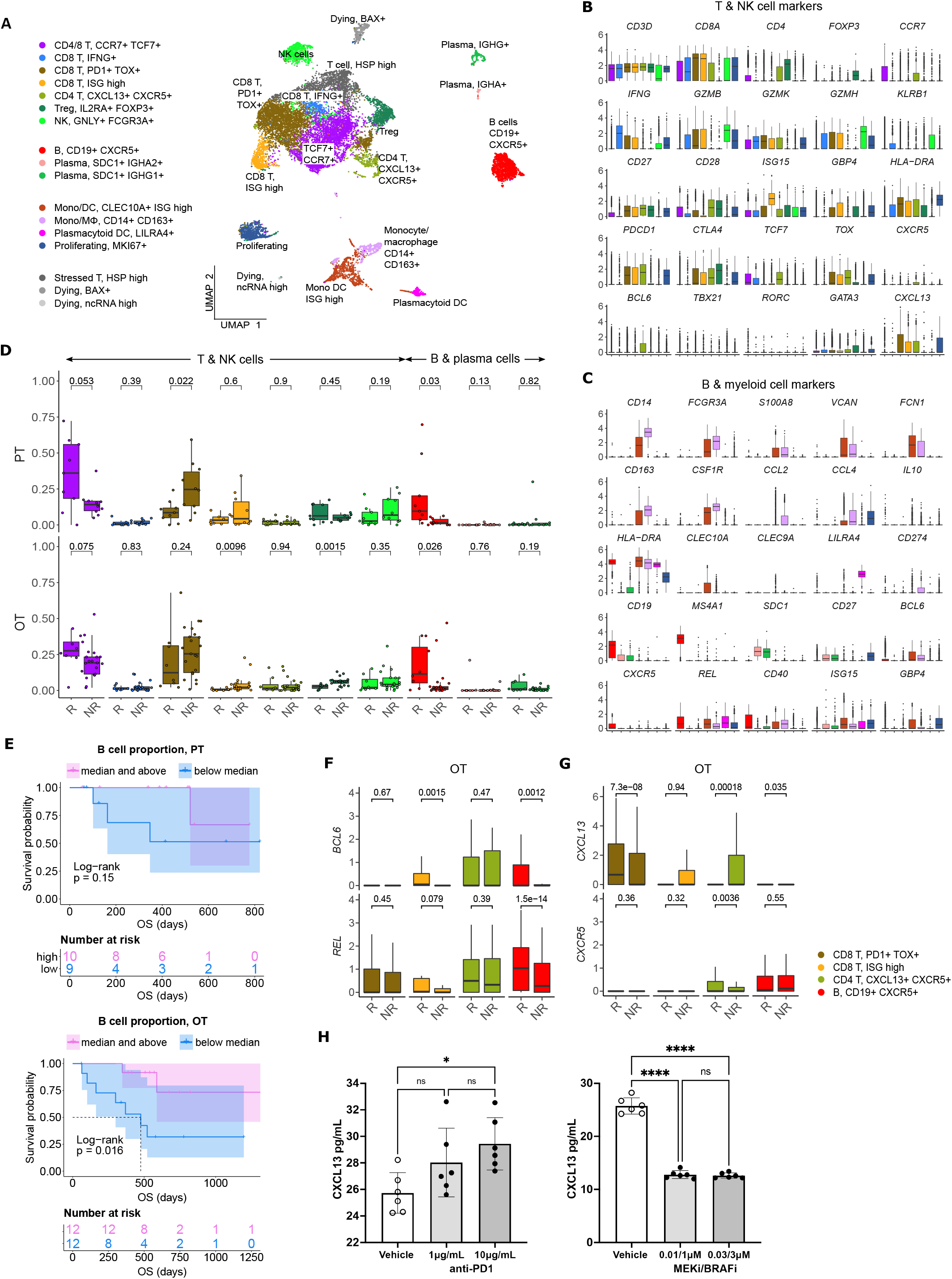
Increased B cell proportion is associated with TLS marker enrichments and improved OS in an independent cohort of ICI treated melanoma. A) UMAP projection of intratumoral CD45+ cells of ICI-treated melanoma. The cell type annotation of each cell cluster was inferred from the DEGs of each cluster. B, C) Normalized expression of markers of cell type/activity among different T and NK cell (B) or B cell and myeloid cell populations (C). D) The fraction of the T, NK B and plasma cell populations in stratified by response vs. resistance to ICI in PT (top) and OT tumors (bottom). The boxplots are colored to match the cell clusters in (A). E) Kaplan-Meier survival curves of ICI-treated patients stratified by the proportion of B cells within either their PT (top) or OT tumors (bottom). F, G) Normalized expression of germinal center (GC) B cell markers, *BCL6* and *REL*, or follicular helper T cells (Tfh), *CXCR5* and *CXCL13*, within the listed T/B cell populations from the scRNA-seq. H) CXCL13 secretion by human peripheral blood mononuclear cells is increased by anti-human programmed death protein (PD-1) antibody treatment but decreased by trametinib (MEK inhibitor) + dabrafenib (BRAF V600E inhibitor) *in vitro*. Significance of pairwise comparisons was computed using t-test, * p<0.05, ****p<0.0001.

T cell-related gene sets were enriched in OT-R groups for both ICI and MAPKi therapies (Fig 1C, Supp. Table 1D). The levels of general T cell marker (*CD3D*), cytotoxic CD8 T cells (*CD8B*) and T cell signature were significantly higher in the OT-R tumors compared to PT and OT-NR tumors for both therapies (Fig 1D, Supp. Fig 1B, Supp. Table 1E). Batch-corrected and normalized T cell marker expressions were not significantly different between ICI and MAPKi in either PT, OT-R or OT-NR samples (Supp. Fig 1C), suggesting minimal, if any, residual batch effects associated with the different datasets. As with T cell markers, increased of IFNγ pathway scores were observed in response to either therapy, however, the scores were not different between responders to ICI and MAPKi (Fig 1E, Supp. Fig 1D, Supp. Table 1F).

Increased T cell infiltration and activities is important for both ICI and MAPKi response (*13*, *15*, *17*, *22*, *32*, *33*, *37*). Intriguingly, only higher normalized expression of CD8 T cell marker *CD8B* or higher IFNγ gene set enrichment in ICI OT tumors, but not MAPKi OT tumors, was significantly correlated with longer survival (Fig 1F). Neither *CD8B* expression nor IFNγ pathway score in PT biopsies was significantly correlated with survival (Supp. Fig 1E), indicating the importance of T cell infiltration and activity after therapy. There was one caveat: the MAPKi RNA-seq datasets only provided progression free survival (PFS) information. Nonetheless, we confirmed that melanoma patients’ PFS is significantly correlated with their OS in a separate microarray datasets of MAPKi treated tumors with both PFS and OS information (*38*, *39*) (Supp. Fig 1F, Supp. Table 2). As such, we use PFS differences in the MAPKi cohort as a surrogate of OS differences.

We noted higher total expression of T cell cytotoxicity-related genes *GZMB*, and *PRF1* in ICI OT-R than MAPKi OT-R (Fig 1G; *PRF1* is higher in ICI OT-R but not statistically significant). We further estimated the level of these cytotoxicity-related genes on a per T cell basis by computing the ratio between normalized expression of the markers and *CD8B*. On this approximated per-CD8 basis, MAPKi-treated responders had lower *GZMB* and *PRF1* expression than their patient-matched PT tumors while ICI-treated responders did not show such decrease (Fig 1H). Overall, our analysis demonstrated more cytotoxic CD8 T cells after ICI treatment as compared to MAPKi, which may explain the stronger correlation between patient OS after ICI therapy and the amount of tumor infiltrating CD8 T cells or their IFNγ signaling pathway activity.

### B cells were more abundant in ICI OT tumors and were correlated with improved survival after ICI therapy

We next examined ICI-specific DEGs to discover additional cell populations or pathways that are significantly associated with response to ICI therapy. Genes related to CD19+ B cells (Fig 31C) and B cells and DC gene sets from (Fig 2A, Supp. Table 1D) were upregulated in ICI OT-R tumors compared to MAPKi OT-R tumors (after adjusting against the average expression of the same genes in the respective OT-NR groups, see Methods). On the other hand, MAPKi OT-R tumors showed enrichment of non-immune related gene sets such as neuron, melanocyte and adipocyte related genes (Supp. Fig 2A, Supp. Table 1D).

Upregulations of B cell gene markers, B cell signature and immunoglobulin genes in OT-R tumors, with respect to either OT-NR or PT tumors were higher after ICI compared to MAPKi therapy (Fig 2B, Supp. Fig 2B, Supp. Table 1E). Importantly, the expression of TLS-associated, germinal center B cell (GC B cells) and follicular helper T cell (Tfh) markers, *CXCR5* and *BCL6*, were significantly upregulated only in response to ICI but not MAPKi (Fig 2C). The presence of tertiary lymphoid structures (TLS) has been correlated with improved survival after ICI therapy in melanoma (*28*, *29*) and other cancers (*30*, *31*). ICI treatment also induced a more significant increase of gene set scores of two TLS signatures (*36*) compared to MAPKi (Supp. Fig 2C, Supp. Table 1F). Among ICI-specific differentially expressed of cytokines and chemokines was *CXCL13*, which is the ligand for *CXCR5* (Fig 2D). The upregulation of *CXCL13* after ICI treatment is expected to recruit *CXCR5*+ B cells and *CXCR5*+ Tfh cells. Another ICI-specific chemokine was *CCL21*, which is expressed by stromal cells of activated lymphoid follicles in secondary lymphoid organs (SLO) and TLS (*40*, *41*).

The magnitude of *CD19* expression and TLS signature score both significantly correlated with survival after ICI therapy but not MAPKi (Fig 2E). On the other hand, the level of B cell marker or the TLS signature in the PT samples is not associated with survival after ICI treatment (Supp. Fig 2D). In agreement with the survival data, B cell marker expression was positively correlated with tumor response to ICI in OT but not PT tumors (Supp. Fig 2E), suggesting level of B cell and TLS marker genes pre-therapy are relatively weak predictors of response and survival after ICI treatment. Thus, response to ICI is marked with increased expression of B cell markers, accompanied with markers (*CXCR5*, *BCL6*), gene signatures and secreted factors (*CXCL13*, *CCL21*) that are indicative of TLS presence in ICI OT-R tumors.

### Enrichment of B cell and TLS gene markers in single cell transcriptome of ICI responding melanoma

To analyze potential connections between enhanced CD8 T cell cytotoxicity and increased presence of TLS-associated B cells in ICI OT-R tumors, we examined single cell transcriptomic data of tumors from an independent cohort of ICI-treated melanoma patients (*33*). Since this scRNA-seq is done on a sampling of sorted CD45+ cells, we can only compare the relative abundances of immune populations within the samples. Nonetheless, the single cell characterization of the immune cells will still allow us to identify significantly increased/decreased expression of gene markers within each immune population.

After normalization and scaling of the gene expression values (see Methods), we re-clustered the single cells and identified the cell types based on differentially upregulated immune gene markers in each cluster (Fig 3A, B, C, Supp. Table 3A). Clusters of known immune populations, such as memory T cells (*CD4/8*+ *CCR7*+ *TCF7*+), activated CD8 T cell (*IFNG+* or high in interferon downstream genes (ISGs)), activated/exhausted CD8 T cell (*PDCD1*+ *CTLA4*+ *TOX*+) with high expression of *CXCL13*, B cell (*CD19*+ *MS4A1*+), plasma cell (*SDC1*+ *IGHG/A*+), NK cells (*FCGR3A*+ *GNLY*+), Tregs (*FOXP3*+ *CTLA4*+), monocyte/macrophage (Mφ) (*CD14*+ *CD163*+), monocyte-derived dendritic cells (mono DC: *CD14*+ *CLEC10A*+), plasmacytoid DC (*LILRA4*+) and proliferating cells (*MKI67*+), were marked accordingly. We further identified a subset of *PDCD1*+ *CXCL13*+ *CTLA4*+ *TOX*+ *TCF7*+ *CD4*+ T cells, which also expressed *CXCR5* and *BCL6* (Fig 3B); this potentially is a population of activated *CXCR5*+ *CXCL13*+ follicular helper T cells (Tfh). We also noted the expression of *BCL6* and *REL* in the B cell population (Fig 3C); along with *BCL6*, *REL* is a transcription factor that is upregulated in GC B cells (*42*).

As the scRNA-seq is done on the CD45+ fraction of the tumor (i.e intratumoral immune cells), we could ask whether the proportions (but not absolute numbers) of each cell population were different in the responder vs. non-responder to ICI. Corroborating the B cell marker up-expression in ICI OT-R tumors in bulk RNA-seq, we noted a significantly increased proportion of B cells in ICI OT-R samples compared to OT-NR samples (Fig 3D, red boxplots, bottom, Supp. Table 3B). We also noted that, in this scRNA-seq cohort, the proportion of B cells was already higher in the PT-R samples (Fig 3D, red boxplots, top, Supp. Table 3B). However, only B cell abundance in ICI OT tumors was significantly correlated with the patients’ OS (Fig 3E, Supp. Table 3C).

Looking at the other cell populations, the proportion of *CCR7*+ *TCF7*+ memory T cells per sample were higher in the PT-R and OT-R samples (Fig 3D, leftmost boxplot, Supp. Table 3B), in line with the original publication (*33*). Populations with higher proportion in the OT-NR biopsies are the proliferating cluster and CD14+ CD163+ Mφ clusters (Supp. Table 3A). The proliferative cluster showed high expression of *CD3D*, *CD8A*, *PDCD1*, *CTLA4* and *TOX* (Fig 3B, righmost boxplot); this population matches the phenotype of an intermediate exhausted cytotoxic T cells, which are expected to differentiate into terminally exhausted T cells (*43*). The CD14+ CD163+ Mφ fraction was associated with worse OS after ICI (Supp. Fig 3B), suggesting that this is an immunosuppressive Mφ population.

In each cell population, we analyzed genes increased in expression in ICI OT-R tumors compared to OT-NR tumors (Supp. Table 3D). Consistent with evidence for GC B cells and Tfh in ICI-responders from the bulk RNA-seq cohort above, CXCR5+ B cells in OT-R showed increased expression of *BCL6* and *REL* (Fig 3F). We also confirmed that expression of *CXCR5* was significantly increased in Tfh/Tph population of ICI OT-R patients (Fig 3F, green boxplots), implying a higher proportion of Tfh cells in ICI OT-R tumors. The bulk RNA-seq cohorts showed an increased overall expression of *CXCL13*, which is the chemotactic factors for the CXCR5+ GC B and Tfh cells in ICI OT-R tumors. The scRNA-seq data allowed us to trace the source of *CXCL13* expression; *CXCL13* was significantly upregulated in the *PDCD1*+ *TOX+* CD8 T cell in ICI OT-R tumors (Fig 3G). Of note, *CXCL13*+ *PD1*+ *TOX*+ CD8 T cell population was recently reported to comprise a high fraction of tumor antigen-specific CD8 T cells (*44*). We next tested if ICI can directly increase CXCL13 expression in antigen activated T cells. We cultured *in vitro* donor-derived peripheral blood mononuclear cells (PBMC) with agonistic antibodies of CD3 and CD28 in order activate and expand the T cells. We confirmed that 5 days of PD-1 antibody treatment increased the secretion of CXCL13 protein (Fig 3H, left). Strikingly, combined treatment with BRAF and MEK inhibitors decreased CXCL13 secretion by half (Fig 3H, right).

CXCL chemokine-receptor interaction analysis using CellChat predicted that *CXCL13* expressed by *PD1*+ *TOX*+ CD8 T cells (source) mainly engaged CXCR5+ B cells (target) (Supp. Fig 3C). On the other hand, ICI OT-NR tumors show a mixture of CXCL16-CXCR6 and CXCL13-CXCR3 interaction among the monocytic DC, macrophages and T cells (with weak CXCL13-CXCR5 interaction involving B cells). Although the relative fractions and normalized expression of *CXCL13* and *CXCR5* were similar between ICI OT-R and OT-NR (Fig 3H, dot plots), the CXCL13-CXCR5 interaction is expected to be stronger in ICI OT-R group since there was a larger fraction of CXCR5+ B cells within the CD45+ immune cells of the latter. Finally, AUCell analysis (*45*) using showed enrichments of TLS gene sets specifically in B and Tfh cell clusters (Supp. Fig 3D), indicative of TLS formation within which CD8+ T, CD4+ Tfh and GC B cells are interacting. Overall, ICI-mediated release of T cell checkpoint engagement of tumor-specific, PD1+ TOX+ CD8 T cells upregulates CXCL13, which subsequently recruits CXCR5+ B cells and Tfh to form TLS in the TME.

### B cells and Tfh in ICI-responding tumors upregulate markers of productive antigen presentation

Using their B cell receptor (BCR), B cells can selectively capture antigens and present them to Tfh cells in through MHC II antigen presentation pathway (*46*–*48*). For this interaction to happen, antigen-specific B cell clones must encounter cognate antigen-specific T cells in the T cell zones of the lymphoid follicle of the SLO or TLS. Indeed, our receptor-ligand interaction analysis suggested that intratumoral B cells in ICI OT-R tumors were presenting antigens to *CXCR5*+ Tfh and Tregs through MHC II pathway (Fig 4A). In OT-NR tumors, MHC II interaction was observed mostly between Tregs and multiple MHC II+ populations, including B cells, DC and the immunosuppressive Mφ (Fig 4A). Overall, the most enriched receptor-ligand interactions involving B cells in ICI OT-R melanoma was the MHC class II pathway (Supp. Fig 4A).

**Fig 4:**
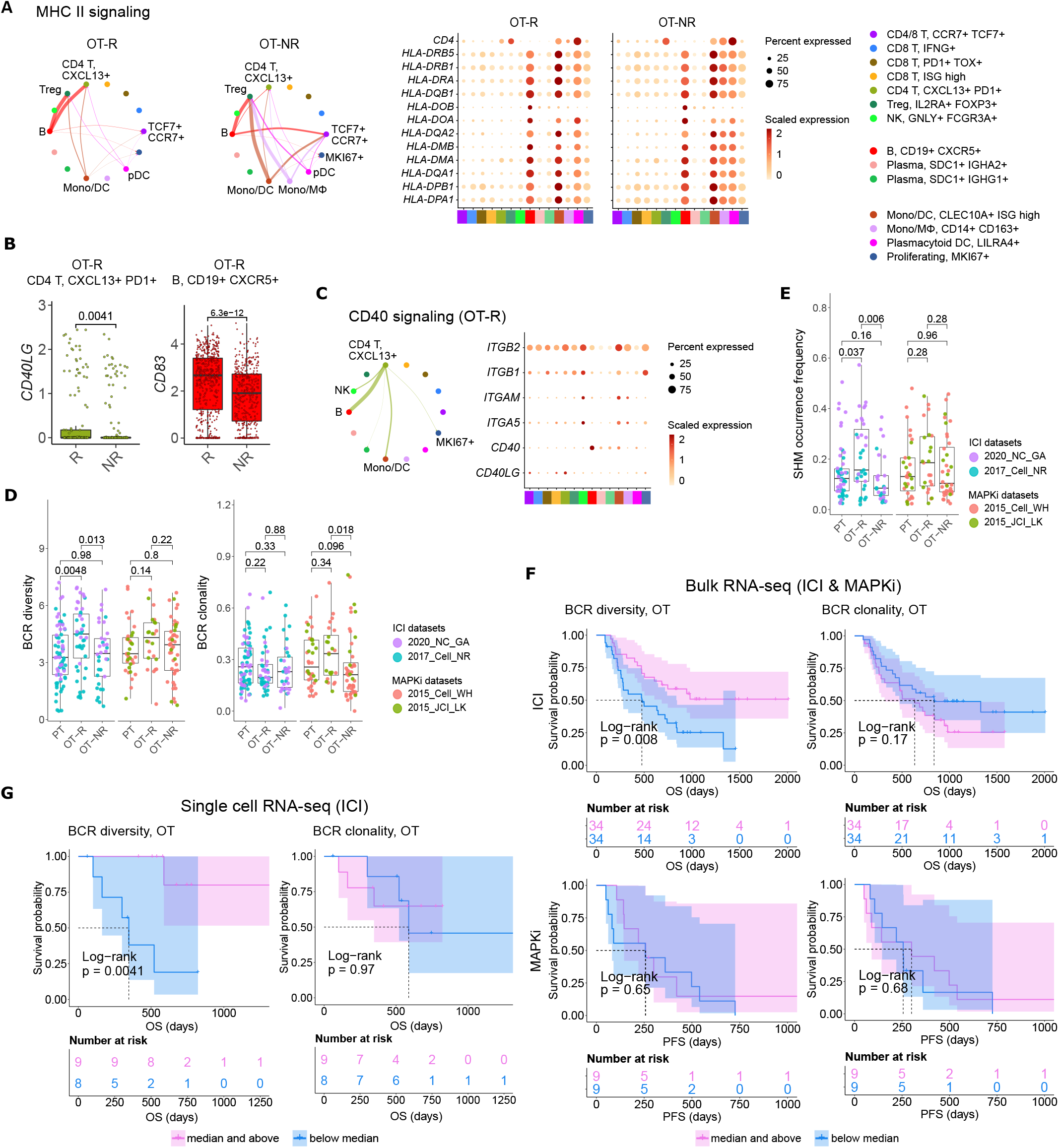
B cells in ICI OT-R tumors present antigens to *PDCD1*+ *TOX*+ tumor-reactive CD8 T and CD4 Tfh cells via the MHC I and II pathways. A) Predicted enrichment of cell-to-cell interaction through MHC II antigen presentation pathway (left) and the normalized expression of MHC II-related genes (right). The expressions of MHC II genes are similar between ICI OT-R and OT-NR but, because of the higher proportion of B cells in ICI OT-R, the predicted MHC II interaction is (relatively) dominated by B cells in ICI OT-R. The interaction is more evenly distributed among the monocytic DC, macrophages, and B cells in ICI OT-NR tumors. B) Normalized expression of *CD40LG* and *CD83* in follicular helper T cell (Tfh) and CXCR5+ B cell populations, respectively. C) Predicted enrichment of cell-to-cell interaction through CD40L/CD40 pathway specific to ICI OT-R tumors (left) and the normalized expression of CD40 pathway related genes (right). D) BCR clonotype diversity and clonality among the PT, OT-R and OT-NR tumors in the ICI or MAPKi therapy group. The BCR clones are based on TRUST4’s predicted CDR3 sequences of the immunoglobulin heavy (IGH) chains in each RNA-seq sample (see Methods). E) Somatic hypermutation (SHM) frequencies based on predicted germline (i.e. prior to SHM) BCR clones by SCOPer (see Methods). The fraction is calculated as the number of germline BCR clones with more than one SHM-generated, mature BCR clones. F) Kaplan-Meier survival curves of ICI- or MAPKi-treated melanoma patients in the bulk RNA-seq datasets stratified by either the BCR diversity (left) or clonality (right) of their OT tumors. G) Kaplan-Meier survival curves of ICI-treated patients in the scRNA-seq dataset stratified either the BCR diversity (left) or clonality (right) of their OT tumors.

Antigen recognition by Tfh results in the activation their T helper function, providing stimulatory signal to CD40+ B cells through upregulation of CD40L (*46*). BCR ligation and CD40/CD40L pathway activation of B cells stimulates expression of APC maturation marker, *CD83*, which further strengthen the antigen presentation activity of the B cells (*49*). Accordingly, we observed the upregulation of *CD40LG* and *CD83* transcripts in the Tfh and B cell populations, respectively (Fig 4B, Supp. Table 3D). Receptor-ligand interaction analysis on the genes in CD40 pathway also showed enrichment of the pathway only in ICI OT-R tumors (Fig 4C, Supp. Fig 4A). Curiously, MHC I pathway interaction ICI OT-R tumors was also dominated by B cells. The latter was predicted to cross-present antigens to *CXCL13*+ *PD1*+ *TOX*+ CD8 T cells (Supp. Fig 4B). CD40L and IFNγ-driven activation of B cells was recently shown to induce antigen cross presentation in B cells (*50*) and both factors are shown to be abundant in the TME of ICI OT-R tumors.

Overall, our single cell transcriptome analysis of ICI-treated melanoma demonstrated upregulation of gene markers and enrichment of pathways associated with a productive antigen presentation by B cells to T cells in the TME of ICI OT-R tumors.

### Diversity but not clonality of B cell population correlates with survival after ICI therapy

After a productive antigen presentation to T cells, antigen specific B cells can undergo class switch recombination (CSR), somatic hypermutation (SHM) and differentiation into long-lived plasma cells or memory B cells (*46*). It is unclear if ICI response in melanoma is associated with the formation of tumor-specific, antibody producing cells, as seen in clear cell renal cell carcinoma (*31*). We did not observe association between the relative abundances of plasma cells and response to ICI in the scRNA-seq cohort (Fig 3B). There were also very few cells expressing *CD19* or *MS4A1* within the *KI67*+ proliferating cell cluster, indicating a rarity of proliferating B cells in this dataset (Fig 3C, based on the levels of the *CD19* or *MS4A1* in the rightmost proliferating cell cluster).

We applied TRUST4 to reconstruct the CDR3 regions within mRNA transcripts of immunoglobulin heavy (IGHA/M/G) chain in the bulk RNA-seq and scRNA-seq data (*51*). By defining each distinct IGH CDR3 sequence as a B cell clone, we can obtain an estimate of the B cells’ clonal dynamics after ICI or MAPKi therapy. Response to ICI exhibited statistically significant increase of BCR diversity in ICI OT-R tumors (with respect to both PT and OT-NR) but not in MAPKi OT-R tumors (Fig 4D, Supp. Table 1G). SCOPer (*52*) analysis predicted an increased somatic hypermutation (SHM) in ICI OT-R tumors, which may have contributed to the increase of BCR diversity in ICI OT-R tumors (Fig 4E). The trend is reversed with BCR clonality where ICI OT-R tumors did not display a higher BCR clonality than ICI PT or OT-NR tumors while MAPKi OT-R tumors did (Fig 4D, Supp. Table 1G).

We observed that most B cell clones in the ICI OT-R tumors were newly infiltrating clones after the therapy (Supp. Fig 4C). The same predominance of OT tumor specific clones can also be observed in the T cell populations (Supp. Fig 4D). The clonal dynamics of both the T and B cells after ICI treatment matches the “T cell clonal replacement” event reported in basal/squamous cell carcinoma patients who responded to ICI (*53*). Higher BCR diversity was associated with improved survival after ICI but not MAPKi therapy (Fig 4F). The significant correlation between BCR diversity and OS after ICI therapy was also confirmed in the scRNA-seq cohort (Fig 4G, Supp. Table 3E, Supp. Table 4A). BCR clonality was not associated with patient survival after either ICI or MAPKi therapy (Fig 4F, G).

The observation of increased BCR diversity in response to ICI therapy suggests that B cells’ role is to present T cells with a broad variety of tumor antigens. Since B cells present tumor antigens to T cells through BCR-specific internalization of extracellular tumor antigens (*48*), a more diverse BCR repertoire will improve the chance of a successful antigen presentation. The strong association between patient survival after ICI and BCR diversity, but not clonality, further implies that ICI response depends on successful tumor antigen presentation to T cells and not on the subsequent clonal expansion and differentiation of B cells into long lived antibody producing plasma cells.

### Combined increase in BCR diversity and IFNγ pathway activity correlates with the greatest long-term survival after ICI therapy

Successful antigen recognition by CD8 T cells increased the overall IFNγ expression (Supp. Fig 5A), which subsequently induced IFNγ pathway activation in the TME of ICI OT-R tumors (Fig 1E). IFNγ expression by multiple CD8 T cell populations mostly activated IFNγ pathway in B cell and monocytic DC of ICI OT-R tumors (Fig 5A). The activation of IFNγ pathway in B cells and DC of ICI OT-R tumors is expected further boost their antigen presentation activity, resulting in additional Tfh and CD8 T cell activation. In OT-NR tumors, IFNγ pathway interaction involved mainly the mono DC and the immunosuppressive CD14+ CD163+ Mφ (Fig 5A).

**Fig 5:**
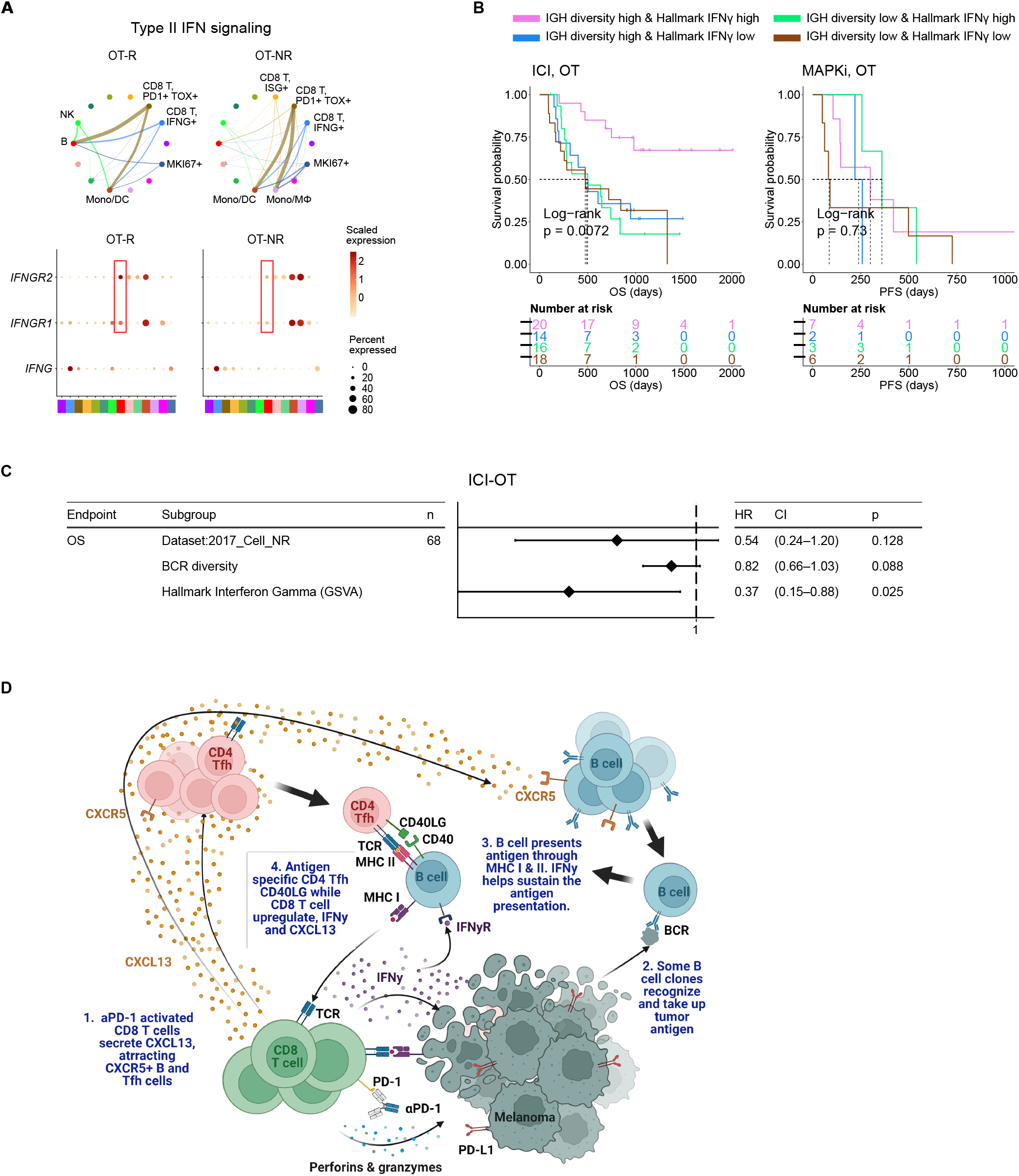
BCR diversity and IFNγ signaling pathway activation are significant factors associated with enhanced OS after ICI therapy. A) Predicted enrichment of receptor-ligand interaction involving type II IFN signaling between *IFNG* expression CD8 T cell clusters and B cells in ICI OT-R tumors (top). Increased expression of *IFNGR1/R2* in the B cells of ICI-responders (bottom) is expected to further enhance B cells’ antigen presentation activity. B) Kaplan-Meier survival curves of patients stratified by BCR diversity and hallmark interferon gamma gene set score in OT tumors of the ICI or MAPKi therapy group. C) Multivariate Cox proportional hazards analysis assessing the hazard ratios of BCR diversity and hallmark interferon gamma gene set score in ICI OT tumor samples. D) The schematic of our proposed model of productive ICI response. First, tumor-reactive CD8 T cells produce CXCL13 in response to ICI to recruit TLS-associated CXCR5+ Tfh and B cells. B cells pick up tumor antigens from tumor cells debris, potentially killed by ICI-activated CD8 T cells. Successful tumor antigen presentation by B cells, expected to highly correlate with their BCR repertoire diversity, results in the activation of the Tfh and (additional) tumor reactive CD8 T cells. Tfh and CD8 T cells upregulates of CD40L and IFNγ, respectively, resulting in enhancement of B cell antigen presentation activity. Finally, newly activated, tumor reactive CD8 T cells kill more tumor cells and secrete more CXCL13 to recruit additional CXCR5+ Tfh and B cells, thus completing an *in situ* cancer immunity cycle.

When stratified by the enrichment of hallmark interferon gamma gene set and BCR diversity of their tumors (high: median and above, and, low: below median), ICI-treated patients with the best survival were those with high hallmark interferon gamma gene set and BCR diversity in their OT samples (Fig 5B, log-rank P=0.0072). Cox proportional hazard analyses suggested that hallmark interferon gamma gene set score and BCR diversity in the OT samples are independent predictors of survival in ICI treated patients (Fig 5C, Supp. Table 4B). Stratification of the patients by the expression of CD8 T cell marker *CD8B* and B cell marker *CD19* show a similar trend where patients with high expression of both *CD19* and *CD8B* had better survival after ICI treatment (Supp. Fig 5B), in agreement with a recent report on a separate cohort of melanoma patients (*29*). Curiously, our analysis also revealed an interaction/dependency between BCR diversity and hallmark interferon gamma gene set scores in relation with the patients’ OS (Supp. Fig 5C, Supp. Table 4C, HR of interaction = 0.55, P = 0.054). Indeed, the combined positive effect of clonally diverse B cells and a high IFNγ signaling pathway activity (indicating successful antigen presentation to T cells) on patients’ OS is significantly greater the sum of their individual effects (Fig 5B). Thus, higher clonal diversity of intratumoral B cells and IFNγ signaling pathway activity in the TME may act synergistically to drive a durable ICI response.

## Discussion

To discover immune factors and pathways associated with durable response to ICI-based immunotherapy, we analyzed the transcriptomic profiles of melanoma biopsies taken before and after ICI treatment. Unique to our study is the use of transcriptomic profiles of melanoma biopsied pre- and post-MAPKi therapy. Since MAPKi therapy also induces significant immune infiltration yet MAPKi-treated patients are less likely to achieve a durable response than ICI-treated patients, comparing immune infiltration associated with ICI against MAPKi allowed us to separate drivers of durable response from immune “bystanders” in ICI OT-R tumors. Genes that are upregulated in ICI OT-R tumors highlighted enrichment of B cell and Tfh gene markers, strongly hinting the presence of TLS in the tumor. This observation confirms revious studies reporting positive association between TLS and ICI response in multiple cancer histologies (*28*–*31*).

In ICI OT-R tumors,the increase of TLS-associated CXCR5+ B cells and Tfh is correlated with increased mRNA expression of CXCR5’s ligand, *CXCL13*, in a population of activated, tumor reactive CD8 T cells (*44*). We propose a model where an effective response to ICI is marked with CXCL13+ CD8 T cell-driven recruitment of highly diversified, CXCR5+ B cell clones, whose subsequent (tumor) antigen presentation activities induce and sustain the activation of tumor-reactive CD4 Tfh and CD8 T cells (Fig 5D). Notably, our model resembles the cancer-immunity cycle initially proposed by Chen and Mellman (*54*) but with the important distinctions of the cycle happening directly in the TME and CXCR5+ B cells functioning as its major APC.

Importantly, our results suggest that B cells mainly function as APCs in ICI OT-R tumors. First, our cell-cell interaction analysis highlighted CXCR5+ B cells as the dominant antigen presenting cells in ICI OT-R TME, presenting antigens to both tumor-reactive CD8 T cells and CD4 Tfh cells and through MHC I and MHC II pathways, respectively. We observed a concomitant overexpression of *CD40LG* in the Tfh population, reflecting a productive antigen presentation by B cells to Tfh (*47*, *48*). CD40L up-expression in Tfh then activated the CD40 signaling pathway in B cells, as shown by upregulation of *CD83* in the latter (*49*). CD83 is a marker of light zone-specific, antigen presenting GC B cells (*46*) and its expression is crucial for B cell longevity after antigen stimulation (*55*). Overall upregulation of IFNγ expression in ICI OT-R tumors (bulk RNA-seq) was predicted to significantly activate the IFNγ pathway of B cells and DC in ICI OT-R tumors. Notably, simultaneous activation of IFNγ and CD40 signaling pathways in B cells can increase their antigen cross presentation to CD8 T cells (*50*, *56*). Successful cross-presentation to tumor reactive CD8 T cells is expected to drive their cytotoxic activity against the tumor.

Another support of B cells’ APC role in ICI response came from the clonal dynamics of B cells in ICI OT-R tumors. Higher BCR diversity, which is expected to increase the chance of tumor antigen uptake and presentation to T cells, is significantly correlated with longer OS after ICI therapy both in the bulk and single-cell RNA-seq cohorts of melanoma patients. On the other hand, higher BCR clonality was not associated with improved OS after ICI, suggesting that B cells’ subsequent clonal expansion, differentiation to long lived plasma cells/memory B cells or specific antibody production are less correlated with response to ICI. Finally, we demonstrated that BCR diversity and IFNγ signaling pathway scores are significant variables that are correlated with survival after ICI therapy. Patients whose tumors display both higher BCR diversity and IFNγ pathway scores after ICI have significantly longer OS than those with only higher BCR diversity or higher IFNγ pathway scores.

Our study has several limitations. First, it is a correlative, retrospective study of combined cohorts of ICI-treated tumors. To overcome this, we ensured that the most important observations from one dataset are corroborated an independent dataset. For instance, the increased expression of TLS markers in ICI OT-R tumors and the association between BCR diversity/clonality with survival were confirmed in both bulk and scRNA-seq datasets of ICI treated melanoma. We also validated the differential effects of ICI (using a PD-1 antibody) and MAPKi on CXCL13 expression in activated T cells. Another limitation is that the use of RNA-seq to reconstruct the CDR3 regions of the TCR or BCR may have limited sensitivity. Additional studies using direct TCR/BCR sequencing of ICI-treated tumors will be needed to confirm our observations.

Our results demonstrate that an effective immune response to ICI involves activation of tumor reactive CD4 and CD8 T cells by antigen presenting B cells, in the context of TLS in the TME. The next logical question is how to leverage this observation in the clinic. Several studies have attempted direct TLS formation using secreted factors such as, CXCL13 (*57*). However, TLS formation may not be sufficient to ensure B cell and T cell activation and the subsequent antitumor immunity (*58*). Strategies to pre-load tumor antigens on B cells to generate B cell vaccine may be more promising as a combinatorial therapy with ICI. B cell-based tumor antigen vaccine was reported to promote ICI efficacy in animal models of melanoma (*56*), lung cancer (*48*) and GBM (*50*). Thus, there is a pressing need for additional pre-clinical and, ultimately, clinical studies to test the most optimal B cell-vaccine approach that can enhance the rate of durable ICI response.

## Methods

### Datasets used

In order to perform a comparative analysis on transcriptomic response to ICI and MAPKi, we analyzed two batches of immunotherapy data and two batches of targeted therapy data. The two immunotherapy datasets are from Riaz N, et al. (*32*) (named 2017_Cell_NR) and Abril-Rodriguez G, et al. (*16*) (named 2020_NC_GA). The patients of these two cohorts were treated with antibodies against the PD-1 receptor (anti-PD-1) including nivolumab and pembrolizumab. The two targeted therapy datasets are derived from Hugo W, et al. (*23*, *34*) (named 2015_Cell_WH) and from Kwong LN, et al. (*35*) (named 2015_JCI_LK). These patients were treated with either BRAF inhibitor monotherapy or BRAF and MEK inhibitors combination therapy (BRAFi: vemurafenib, dabrafenib, encorafenib; MEKi: cobimetinib, trametinib, binimetinib; one patient was treated with trametinib monotherapy). All samples were classified into three groups: pre-treatment (PT), on-treatment responding (OT-R), and on-treatment non-responding (OT-NR). OT-R is defined by clinical benefit after therapy (CR, PR, or SD by RECIST criteria). OT-NR is defined by no clinical benefit (PD) (Supp. Table 1A). For single cell transcriptome analysis, we utilized the data from Sade-Feldman et al (*33*). The data profiled 16,291 immune cells (CD45+ cells) from 48 tumor samples of melanoma patients treated with ICI. All samples were classified into four groups: pre-treatment responding (PT-R), pre-treatment non-responding (PT-NR), on-treatment responding (OT-R), and on-treatment non-responding (OT-NR). The PFS and OS data of MAPKi treated patients (Supp. Table 2) were extracted from two separate publications; both studies were not incorporated in our combined analysis as they only provide microarray datasets (*38*, *39*).

### Gene expression analysis

The bulk RNA-seq data was re-aligned to hg38 human reference genome using HiSAT2 (v2.1.0), then processed using samtools (v1.10) and picard (v2.25.0). The gene expression count was calculated using htseq-count (v0.11.2). The gene expression was normalized using trimmed mean of M-values (TMM) and converted to count per million (cpm) expression value using edgeR R package (v3.32.1). Batch effect was corrected using removeBatchEffect function in the R package limma (v3.46.0). The normalized expression of each gene (in log_2_ cpm) is provided in Supp. Table 1B.

### Differential gene expression analysis

We first computed the expression change of each gene (in log_2_ FC) between the PT and OT samples of each patient (the OT samples can be OT-R or OT-NR). Thus, each gene will have a list of log_2_ FC values associated with therapy response (OT-R with respect to their respective PT) and resistance/non-response (OT-NR w.r.t their respective PT). Differentially expressed genes (DEGs) between the OT-R and OT-NR groups were defined by:

1. two-fold difference between the arithmetic average of log_2_ FCs in OT-R and arithmetic average of log_2_ FCs in OT-NR (Δlog_2_ FC ≥ 1 where Δlog_2_ FC = mean(log_2_ FC(OT-R)) - mean(log_2_ FC(OT-NR))), and,
2. FDR-adjusted t-test p value < 0.05 between the log_2_ FCs of the OT-R and OT-NR groups.

The sets of DEGs that are upregulated in the OT-R groups of either therapy were shown in Fig 3A. The gene sets/ontologies that are enriched in the ICI-specific, MAPKi-specific and ICI and MAPKi shared DEGs were computed using Enrichr (*36*) (see the **gene set analysis** section). Other than comparing the list of DEGs, we also directly computed the difference of log_2_ FC difference between the OT-R and OT-NR groups across the two therapies. For instance, to find the genes that are upregulated at least two-fold higher in ICI’s responder when compared to MAPKi responders (after adjusting with their respective non-responder groups), we selected genes satisfying:

1. ΔΔFC_ICI-MAPKi_ ≥ 1, where ΔΔFC_ICI-MAPKi_ = Δlog_2_ FC_ICI_ – Δlog_2_ FC_MAPKi_, and,
2. Δlog_2_ FC_ICI_ ≥ 1

Conversely, the genes that are upregulated in MAPKi’s responders were computed in the same manner.

### Gene sets analysis

Shortlisted genes from DEG analysis were analyzed for overlap-based gene set enrichment using Enrichr (*36*). Top enriched gene sets from Human_Gene_Atlas and HuBMAP_ASCT_plus_B_ augmented_w_RNAseq_Coexpression collections are visualized in Fig 1C and listed in Supp. Table 1D. Score based, single sample gene set enrichments was computed using Gene Set Variation Analysis (GSVA) (*59*) through R packages GSVA (v1.38.2), GSVAdata (v1.26.0), and GSEABase (v1.52.1).

We used the interferon gene sets from the Molecular Signatures Database (MSigDB) (*60*, *61*). Specifically, we collected gene sets containing the keyword “IFN” or “interferon” from the [H: hallmark gene sets](https://www.gsea-msigdb.org/gsea/msigdb/download_file.jsp?filePath=/msigdb/release/7.0/h.all.v7.0.symbols.gmt), C2 [CGP: chemical and genetic perturbations](https://www.gsea-msigdb.org/gsea/msigdb/download_file.jsp?filePath=/msigdb/release/7.0/c2.cgp.v7.0.symbols.gmt), [C6: oncogenic signatures](https://www.gsea-msigdb.org/gsea/msigdb/download_file.jsp?filePath=/msigdb/release/7.0/c6.all.v7.0.symbols.gmt), and [C7: immunologic signatures](https://www.gsea-msigdb.org/gsea/msigdb/download_file.jsp?filePath=/msigdb/release/7.0/c7.all.v7.0.symbols.gmt) (Supp. Table 1F). The gene set of TLS signatures were manually curated in gmt file format. Cabrita *et al* developed two gene sets that reflected the presence of TLS in melanoma. One TLS signature of nine genes (*CD79B*, *CD1D*, *CCR6*, *LAT*, *SKAP1*, *CETP*, *EIF1AY*, *RBP5*, *PTGDS*) were found using differential expression analysis. The other TLS signature of seven genes (*CCL19*, *CCL21*, *CXCL13*, *CCR7*, *CXCR5*, *SELL*, *LAMP3*) were constructed from a compendium of TLS hallmark genes (*29*).

### Clonotype analysis

As for the TCR clonotyping, the raw output of clonotypes was derived from the raw FASTQ reads of the bulk RNA-seq data using TRUST4 (v1.0.4) with default settings. Nonproductive CDR3aa were removed the raw output. Clonotypes are separated by chain names such as TRA, TRB, TRD, TRG, IGH, IGK, IGL. Convergent clonotypes, which possess the same amino acid sequences but different nucleotide sequences, were merged. As for the BCR clonotyping, considering the SHM mechanism of GC B cells, productive IGH chains were selected from the raw AIRR standard format output derived from the bulk RNA-seq data using TRUST4 (v1.0.4). Then the hierarchicalClones() function in the R package SCOPer (*52*) (v1.2.0) was used to infer—based on the nucleotide sequence—the germline BCR clones that arise from V(D)J recombination (in the bone marrow) and mature (SHM-generated) BCR clones that are derived from germline clones after somatic hypermutation process in the germinal center (*42*). The R package SHazaM (*62*) (v1.1.0) was used to automatically calculate the threshold for the bimodal distribution in the hierarchicalClones() function as defined by the SCOPer pipeline. Consequently, the SHM occurrence frequency can be calculated using the count of germline clones associated with more than one mature BCR clone divided by total count of the germline clones. The clonotype repertoire metrics of TCR and BCR, including count, diversity and clonality were calculated using the custom R package [rTCRBCRr](https://cran.r-project.org/package=rTCRBCRr) (v0.1.0) and other custom R scripts.

### Cell population abundance estimation

The R package MCPcounter (v1.2.0) was applied to the normalized log_2_ cpm expression matrix from the bulk RNA-seq data in order to estimate the absolute abundance of eight immune and two stromal cell populations in each sample, including T cells, CD8 T cells, cytotoxic lymphocytes, B lineage, NK cells, monocytic lineage, myeloid dendritic cells, neutrophils, endothelial cells, and fibroblasts.

### Overlap-based gene set enrichment

The enrichment of specific biological process or gene ontologies were analyzed using Enrichr, which is implemented in a R package enrichR (v3.0) (*36*). Specifically, lists of DEGs were tested for significant overlaps with pre-curated gene sets within the Enrichr based on fisher exact test. The P values we utilized were the ones adjusted for multiple tests using the FDR method and the cutoff of enriched gene set term is adjusted P value < 0.05.

### Survival analysis

The Kaplan-Meier curves were used to visualize differences in survival between patient groups. Cox proportional hazard (Cox PH) regression was used to assess the effect of single or multiple variables on hazard ratio. These analyses were performed using the R packages survival (v3.2.13) and visualized using survminer (v0.4.9), and survivalAnalysis (v0.3.0). For PFS analysis in the MAPKi-treated cohort, we only included OT tumors biopsied prior to progression (giving a total 25 OT tumors from 19 MAPKi-treated patients).

### Single cell analysis

The single cell data was processed using the R package Seurat (v4.0.2). The raw count data were provided by the authors. We first normalized the raw counts using the NormalizeData() function with normalization.method=“LogNormalize” and scale.factor=10000. 17 clusters were identified at resolution=0.5. Cell types of the clusters were manually annotated based on each cluster’s gene markers, which were computed using FindAllMarkers() (min.pct=0.25 and logfc.threshold=0.585). The R package AUCell (v1.12.0) was used to identify gene set enrichments in the single cell transcriptome. The R package CellChat (v1.1.3) was used to visualize cell-cell communication network, grouped by different signaling pathways, among different cell types.

### Statistical information

All the statistical analysis were performed using R programming language version 4.0.5. Unless otherwise stated, all the statistical tests were two tailed. In all boxplots, including gene expression, GSVA score, and gene expression ratio, P values were calculated using a two-sided Wilcoxon rank sum test. In all boxplots, the median is indicated by the line within the box and the 25th and 75th percentiles indicated by the lower and upper bounds of the box. The upper and lower lines above and below the boxes represent the whiskers. Pearson’s R correlation coefficient was computed using R’s *cor.test* function. The P values of the Pearson R were computed using two-sided t-test as described in the documentation. In the Kaplan-Meier survival curves, the P value is the log-rank test P value. In the Cox PH analysis, the P value shown for each variable in the graph is the result of Wald test.

### In vitro assessment of primary human immune cells

Peripheral blood mononuclear cells (PBMC) from healthy donors were isolated from blood by density gradient centrifugation (Ficoll-Hypaque). PBMC were cultured at 10^6^ cells/well in 12 well plates in complete media with 2ME. Stimulation was provided by anti-CD3/CD28 Dynabeads (Invitrogen), as well as anti-human PD-1 (clone pembrolizumab, BioXcell), MEKi trametinib (LSBiosciences) and BRAFi dabrafenib (BioVision), or vehicle. Supernatants were collected on day 5. Secreted CXCL13 was quantified in supernatants by ELISA (R&D Systems). The concentrations chosen for dabrafenib and trametinib was based on reported maximum plasma concentration in patients, which were 2.4 μM and 0.03 μM, respectively (*63*). To avoid overactivation of the T cells, We chose a lower PD-1 antibody concentrations (1 and 10 μg/mL) than the median C_max_ and C_trough_ of pembrolizumab (89.1 and 27.6μg/mL) (*64*).

## Supporting information

Supp. Tables zip

## Data availability

The bulk RNA-seq data were downloaded from the following online repositories. 2017_Cell_NR (ICI) was from PRJNA356761; 2020_NC_GA (ICI) was from PRJNA578193; 2015_Cell_WH (MAPKi) was from PRJNA273359, PRJNA303170, PRJNA403850; 2015_JCI_LK (MAPKi) was from EGAD00001001306. The single cell RNA-seq data were download from the GSE120575. Instead of the TPM normalized expression values, we started from the raw counts provided by the authors in personal communications. The raw count expression values were included in the source code of this study. All the source codes used in this study were uploaded to the GitHub [repository](https://github.com/sciencepeak/TCR_integrated_project).

## Acknowledgments

This study was funded in part by the NIH/NCI grant (1R01CA236910) and grant from Parker Institute for Cancer Immunotherapy. L.D is supported by grant from Parker Institute for Cancer Immunotherapy at UCLA and a fellowship from National Cancer Center. W.H is supported the NIH/NCI grant (1R01CA236910), the Melanoma Research Alliance, the Margaret E. Early Medical Research Trust Grant, and the Parker Institute for Cancer Immunotherapy at UCLA. We thank Moshe Sade-Feldman and Nir Hacohen for generously providing the raw count expression data of single cell RNA-seq used in their publication. Donor derived PBMCs were obtained from the UCLA/CFAR Virology Core Lab (supported by NIH grant 5P30 AI028697).

## Author contributions

L.D, L.S., Y.Z., W.H. designed the experiments and analyses. L.D. implemented of the overall computational analyses unless specified otherwise. L.S. developed the single cell analysis pipeline. L.D., L.S., M.B. performed single cell analysis. L.N.S, M.G.L performed the *in vitro* experiments. L.D, R.M.P, M.A.S, M.G.L and W.H. wrote the manuscript. All authors reviewed and approved the manuscript.

## Materials and Correspondence

Correspondence and requests for materials should be addressed to

## Additional information

Supplementary information is available for this paper online

## Supplementary figures

**Supp. Fig 1:**
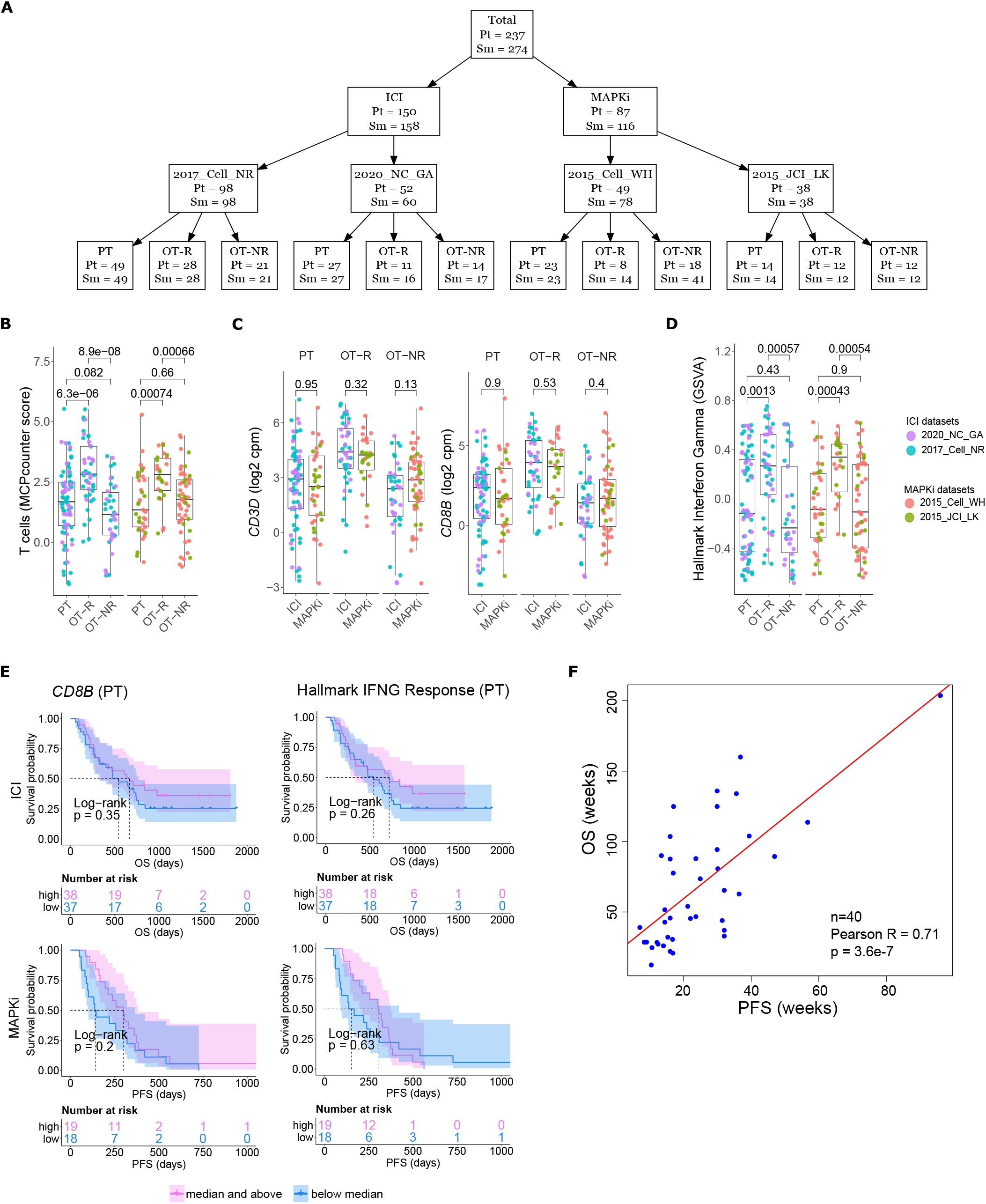
Shared and therapy-specific transcriptomic changes after ICI and MAPKi therapy. A) Schematic of the bulk RNA-seq data sets of ICI- and MAPKi-treated melanoma used in this study. B) T cell enrichment scores computed by MCPcounter in the PT, OT-R and OT-NR samples of patients under ICI and MAPKi therapy. C) Normalized expression of T cell marker genes *CD3D* and *CD8B* in the PT, OT-R and OT-NR samples of patients under ICI and MAPKi therapy (inter therapy comparison). D) GSVA gene set enrichment scores of hallmark interferon gamma gene set from the Molecular Signature database. E) Kaplan-Meier survival curves of ICI- or MAPKi-treated melanoma patients stratified by either *CD8B* expression (left) or hallmark interferon gamma response gene set scores (right) in their PT tumors. F) Correlation between PFS and OS after MAPKi therapy across two separate melanoma microarray datasets.

**Supp. Fig 2:**
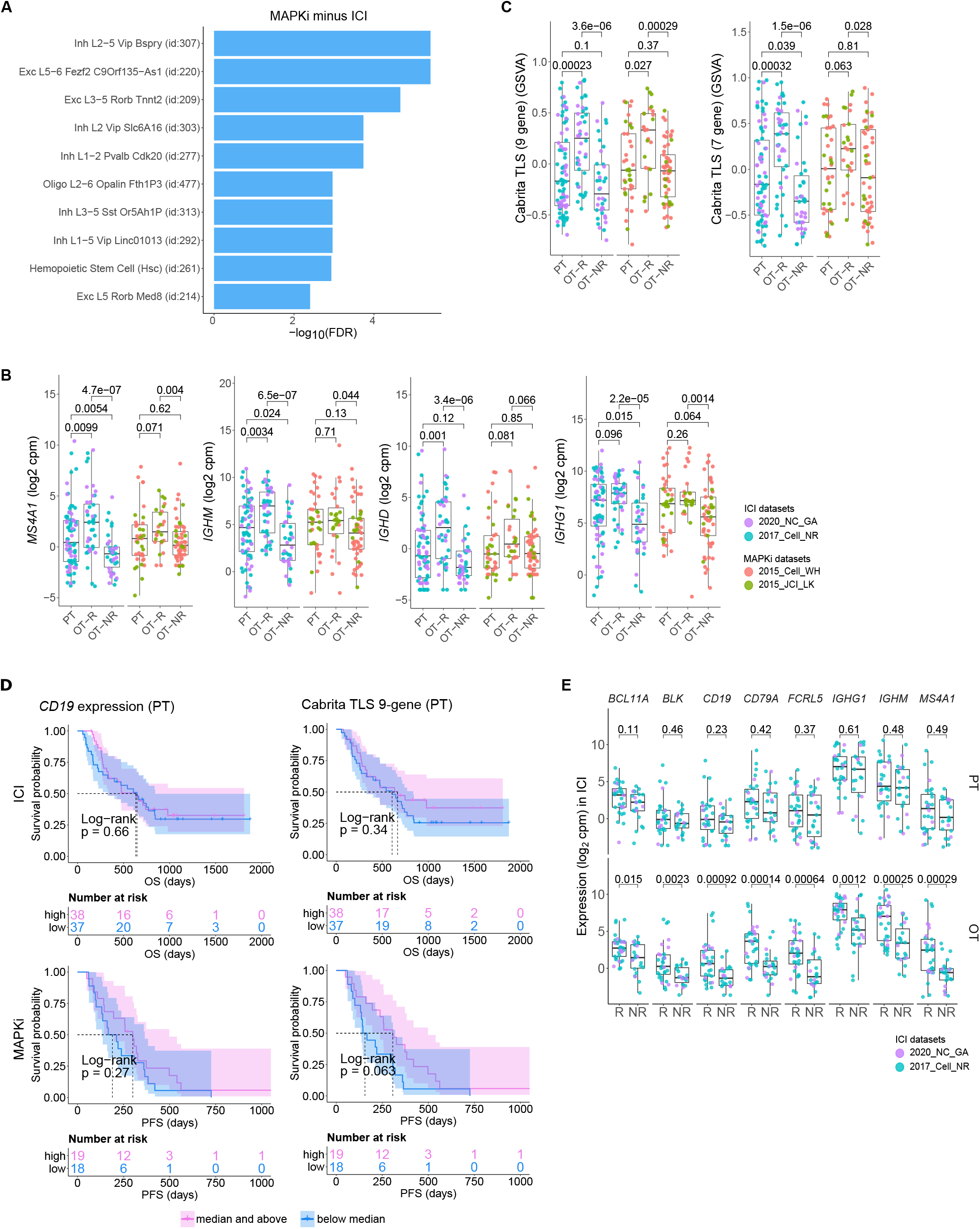
Characterization of B cell and TLS-related gene expression changes after ICI and MAPKi therapy. A) Enriched tissue specific gene sets in DEGs upregulated in MAPKi OT-R tumors with respect to ICI OT-R tumors (after adjustment by respective therapy group’s OT-NR tumors). B) Normalized expression of the listed B cell-related genes among the PT, OT-R and OT-NR tumors in the ICI or MAPKi therapy group. C) Enrichment scores of a recently published TLS 7-gene and 9-gene gene sets among the PT, OT-R and OT-NR tumors in the ICI or MAPKi therapy group. D) Kaplan-Meier survival curves of ICI- or MAPKi-treated melanoma patients stratified by either *CD19* expression (left) or TLS gene set enrichment score (right) in their PT tumors. E) (top) Pairwise expression comparison of the listed B cell-related genes between the PT samples of melanoma patients who will subsequently respond (R) or not respond (NR) to ICI. (bottom) The same comparison applied on ICI OT tumors.

**Supp. Fig 3:**
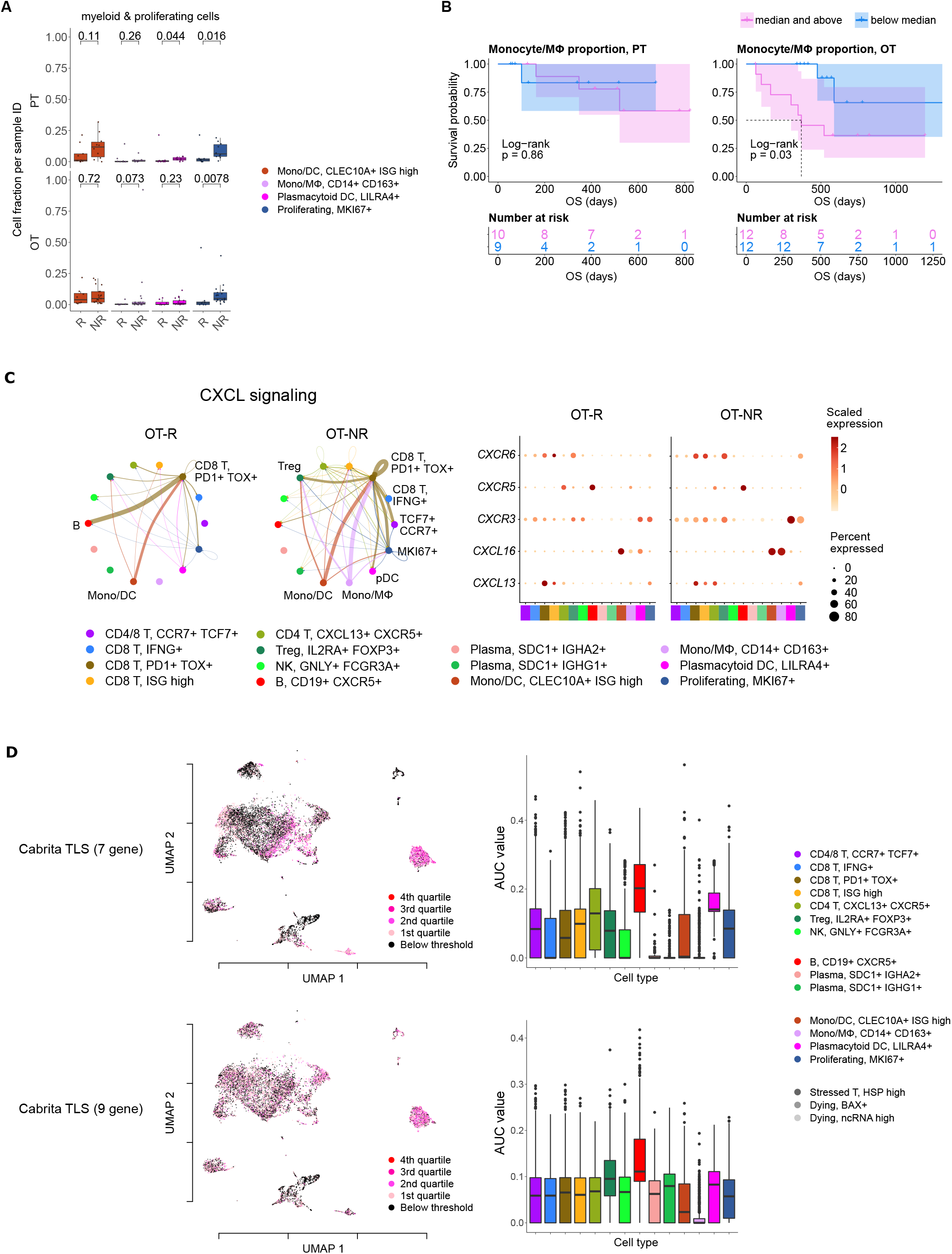
Single cell analysis of ICI-treated melanoma. A) The fraction of the myeloid and proliferating cell populations in stratified by response vs. resistance to ICI in PT (top) and OT tumors (bottom). B) Kaplan-Meier survival curves of ICI-treated patients stratified by the proportion of monocyte/monocyte-derived macrophages within either their PT (top) or OT tumors (bottom). C) Predicted enrichment of receptor-ligand interaction involving the CXCL chemokine signaling in the ICI OT samples (left; the color of the connecting edge is the source cell’s) and the normalized expression of the chemokine and their receptors in each immune population (right). The interaction involving the CXCL13-CXCR5 pair between tumor reactive, *PDCD1*+ *TOX*+ CD8 T cells and B cells was more enriched in ICI OT-R tumors. D) Single cell-based gene set enrichment score of the TLS 7-gene and 9-gene signatures projected on the UMAP (left) or presented in boxplot across all cell types (right). The scores were separated into quartiles to assist the visualization of high and low gene set enrichments in different cell populations.

**Supp. Fig 4:**
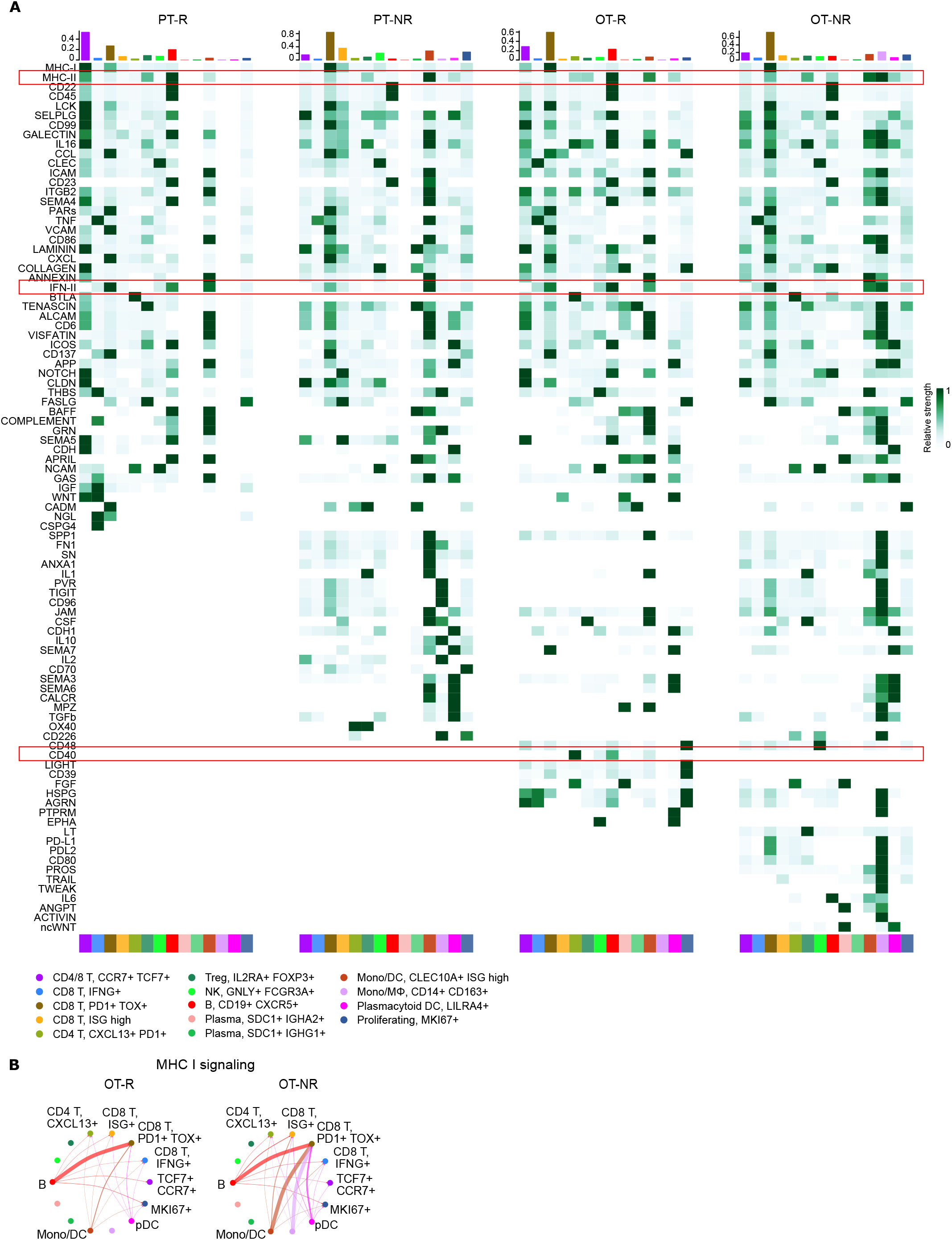
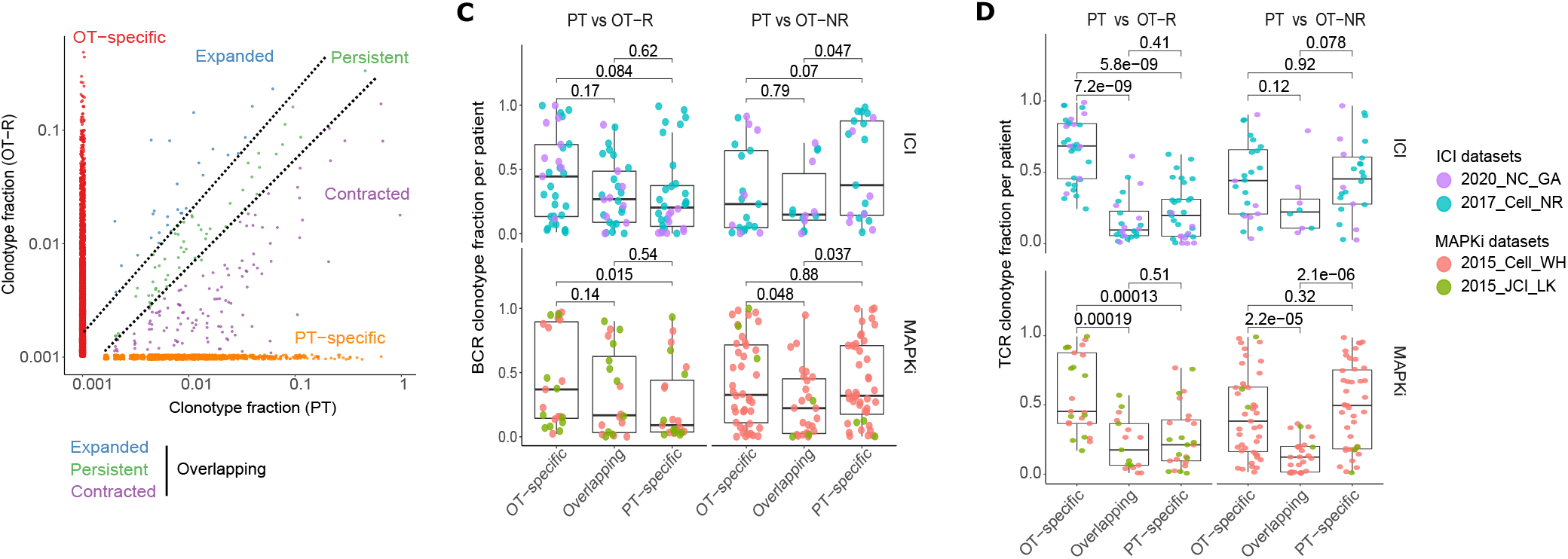
B cell-associated antigen presentation and clonotype analyses. A) Inferred cell-cell communication among intratumoral immune populations of pre- and post-ICI treated melanoma across curated signaling pathways in CellChat. The interactions are grouped based on response vs. no-response to ICI in either PT or OT tumors. B) Predicted enrichment of cell-to-cell interaction through MHC I antigen presentation pathway. C, D) Change in BCR (C) or TCR (D) clonal fraction in grouped by clones found only in the OT sample (OT-specific), only in the PT (PT-specific) and both in the PT and OT samples (overlapping, see illustration on the left). These fractions are calculated with respect to the union of all BCR/TCR clones found in the PT and OT samples of each patient; this analysis is done only on patients with PT and OT tumors.

**Supp. Fig 5:**
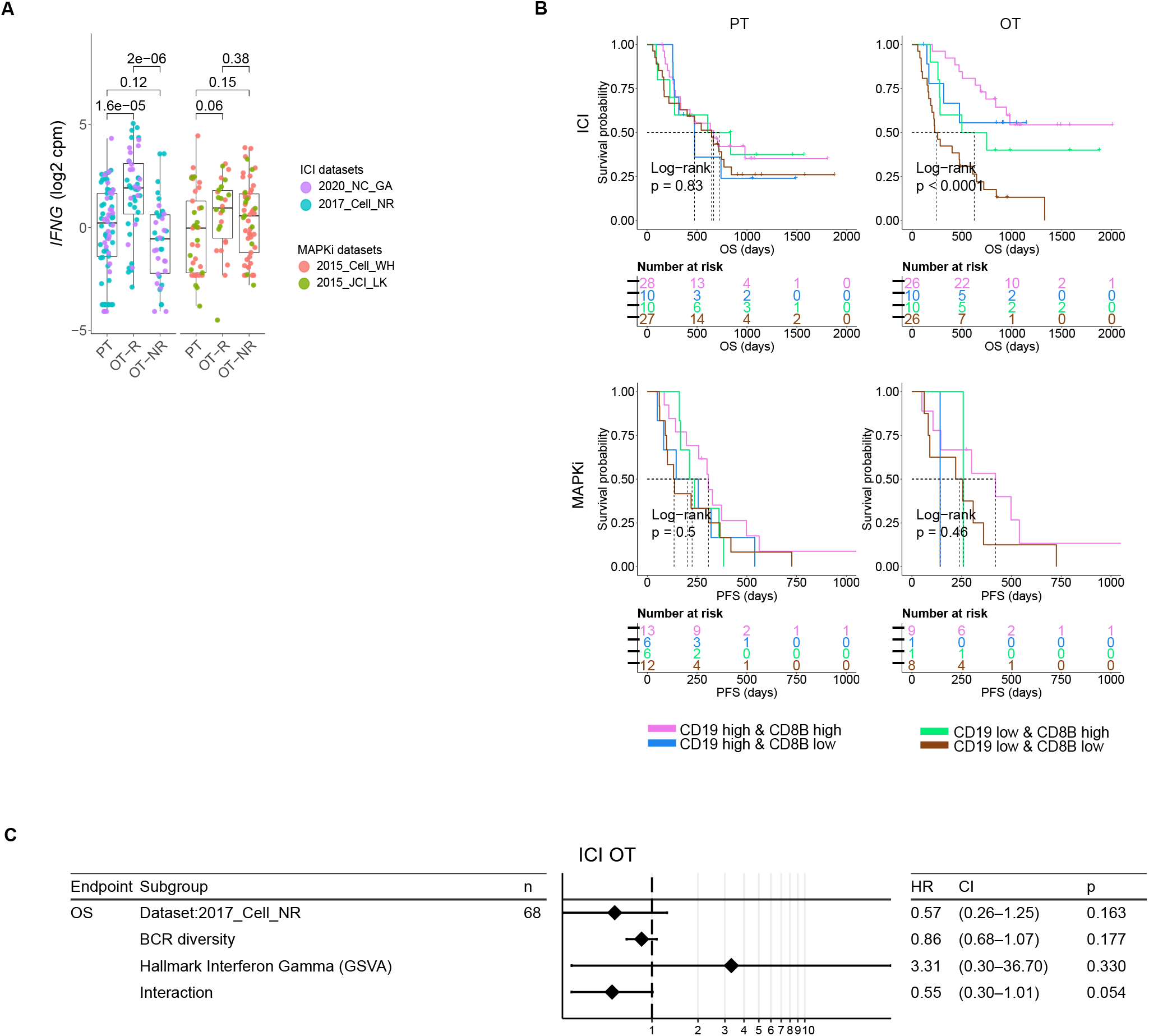
Multivariate analysis of survival examining the association among T cell and B cell related variables to patient OS after ICI therapy. A) Normalized bulk RNA-seq expression of *IFNG* in the PT, OT-R and OT-NR samples of patients under ICI and MAPKi therapy. B) Kaplan-Meier survival curves of patients stratified by normalized expressions of *CD8B* and *CD19* in either PT or OT tumors of the ICI or MAPKi therapy group. C) Multivariate Cox proportional hazards analysis assessing the hazard ratios of BCR diversity, hallmark interferon gamma gene set score, and their interaction in ICI OT tumor samples.

## Supplementary data

Supp. Table 1: Clinical characteristics, gene expression and gene set enrichments in the bulk RNA-seq datasets.

A) Summary of clinical data associated with the tumor samples (bulk and scRNA-seq datasets).

B) Normalized expression matrix (log_2_ cpm, batch corrected) of the bulk RNA-seq data of ICI or MAPKi-treated tumors.

C) Difference in fold changes of OT-R vs. OT-NR samples after ICI or MAPKi therapy.

D) Gene sets/cell markers enriched in tumors responding to ICI, MAPKi or both (Enrichr analysis).

E) The relative abundance of immune and stromal cell populations in each sample in bulk RNA-seq dataset (MCPcounter analysis).

F) GSVA score matrix of selected gene sets of the bulk RNA-seq data of ICI or MAPKi-treated tumors.

G) TCR and BCR clonotype repertoire metrics of the ICI and MAPKi datasets (bulk and scRNA-seq).

Supp. Table 2: PFS and OS correlation in MAPKi-treated melanoma patients.

A) OS and PFS data from two MAPKi treated gene expression microarray datasets (Rizos et al CCR 2014 and Long et al Nat. Comm. 2014).

Supp. Table 3: Cell type fraction, DEGs and survival analysis of scRNA-Seq data set of ICI-treated melanoma.

A) Differentially upregulated genes in each single cell cluster of CD45+ cells from ICI-treated tumors.

B) The fraction of immune cell populations in each PT/OT tumor sample of ICI-treated melanoma patients.

C) The association between the fraction each immune population (in PT or OT tumors) and OS after ICI therapy.

D) Differentially expressed genes in PT and OT tumors based on response (R) vs. no-response (NR) to ICI.

E) The association between TCR/BCR clonality or diversity in PT or OT tumors and OS after ICI therapy.

Supp. Table 4: Univariate and multivariate survival analyses of B cell, T cell and TLS-related gene/gene sets.

A) Univariate CoxPH analysis of survival of B cell, T cell and TLS-related gene/gene sets (bulk RNA-seq cohorts).

B) Multivariate CoxPH analysis of survival of significant B cell, T cell and TLS-related gene/gene sets (ICI-OT only, pairwise independent).

C) Multivariate CoxPH analysis of survival of significant B cell, T cell and TLS-related gene/gene sets (ICI-OT only, with pairwise interaction term).

